# MICROBIOME-MODIFIED METABOLITES PREDICT ACUTE BRAIN INJURY OUTCOME

**DOI:** 10.1101/2024.06.24.599530

**Authors:** Orlando DeLeon, Hugo D Ceccato, Ashley Sidebottom, Nada Babtain, Sara Roy, Jason Koval, William H Roth, David G. Beiser, Ali Mansour, Fernando D Goldenberg, Eugene B Chang, Christos Lazaridis

## Abstract

**BACKGROUND:** Acute brain injury (ABI)-mediated disruption of the microbiome may potentiate inflammation and secondary brain injury (SBI). However, microbial-specific mediators and mechanisms remain unclear.

**METHODS:** Thirty-five consecutive patients with ABI admitted to the neuroscience critical care unit at the University of Chicago were prospectively studied. Injury severity at hospital admission was assessed using the Injury Severity Score (ISS) and the Glasgow Coma Scale (GCS). Final neurologic function was assessed via the Glasgow Outcome Score extended (GOSe). Serum, plasma, and stool targeted metabolomics, as well as stool shotgun metagenomics, were performed on longitudinal samples collected during hospitalization.

**RESULTS:** Multivariate analysis identified microbiome-modified metabolites that were positively and negatively associated with functional outcomes after ABI. Novel identification of conjugated bile acid (BA) species and vitamin B12 precursors indicative of outcome were detected in the first collected samples (within 48 hours). Network analysis revealed greater integration of negatively associated metabolites across tissues and identified tauro-α/μ-muricholic acid (TMCA) as central to the cross-tissue metabolomes.

**CONCLUSIONS:** Microbiome metabolites may be useful in assessing brain injury outcomes to inform treatment. Bile acid species transformed by the gut microbiome are predictive of ABI outcome.

## Introduction

Stroke and traumatic brain injury (TBI) are leading causes of death and disability.^1^ Acute brain injury (ABI) after TBI or a cerebrovascular event (intracranial hemorrhage or acute ischemic stroke) is followed by a series of cellular and molecular pathobiological changes that collectively contribute to secondary brain injury (SBI).^2,3^ The focus of clinical care after ABI is to prevent, detect, and mitigate SBI that causes additional irreversible brain damage and contributes to poor patient outcomes.^4^

Preclinical models have demonstrated that ABI disrupts intestinal motility and permeability, leading to gut microbiome composition perturbations and a host-maladaptive state known as gut dysbiosis.^5^ Gut dysbiosis also affects the integrity of the blood brain barrier (BBB).^6^ Coupled with ABI-induced disruptions to the BBB, intestinal molecules and pro-inflammatory immune responses more easily permeate the central nervous system (CNS), resulting in increased microglial activity, neuroinflammation, and neuropathology.^7^

Preclinical models have also demonstrated that stroke induces changes to gut microbiota composition, an early loss of gut integrity, bacterial translocation into the host, and alterations in microbial function.^8^ These studies suggest the existence of a complex microbiota-gut-brain axis (MGBA), revealing microbiota at the time of injury influence the degree of neuropathology and functional impairment following ABI. Monitoring the nature and extent of gut-dysbiosis may provide a diagnostic tool for ABI phenotyping and treatment guidance. Furthermore, perturbations in bacterial composition appear 24–72h following brain injury corresponding to SBI pathophysiology, representing a clinically-relevant treatment window.

Alterations in gut microbiota have been also observed in humans with ABI.^9,10^ Compositional profiles are distinct from other neurological diseases suggesting ABI-specific microbiome mechanisms.^11^ While inter-personal and intra-population microbiota variation has confounded progress, microbially produced or modified small molecules are highly conserved across individuals and species, prompting increased use of metabolomics in evaluating microbiota function in disease states. Many of these molecules transducing effects on health. For example, host-produced bile acids (BAs) modified by gut microbiota and reabsorbed into the systemic circulation exert immunomodulatory properties during TBI^10^, with serum BA levels reduced in TBI subjects.^12^ Recently, a human study demonstrated metabolite associations with TBI outcome but did not specifically examine microbiome-specific metabolites.^13^ Therefore, we undertook a combined cross-sectional and longitudinal study to specifically examine microbial metabolites and associations with ABI outcome.

## Methods

### Study Participants

Thirty-five consecutive patients with ABI admitted to the neuroscience critical care unit (NCCU) were prospectively studied at the University of Chicago (Fig. 1). Severity of injury at hospital admission was assessed with the Injury Severity Score (ISS) and Glasgow Coma Score (GCS) with outcome of neurological function assessed at discharge by the Glasgow Outcome Score extended (GOSe) (Table 1 and Supplementary Table 1).

**Figure 1.**
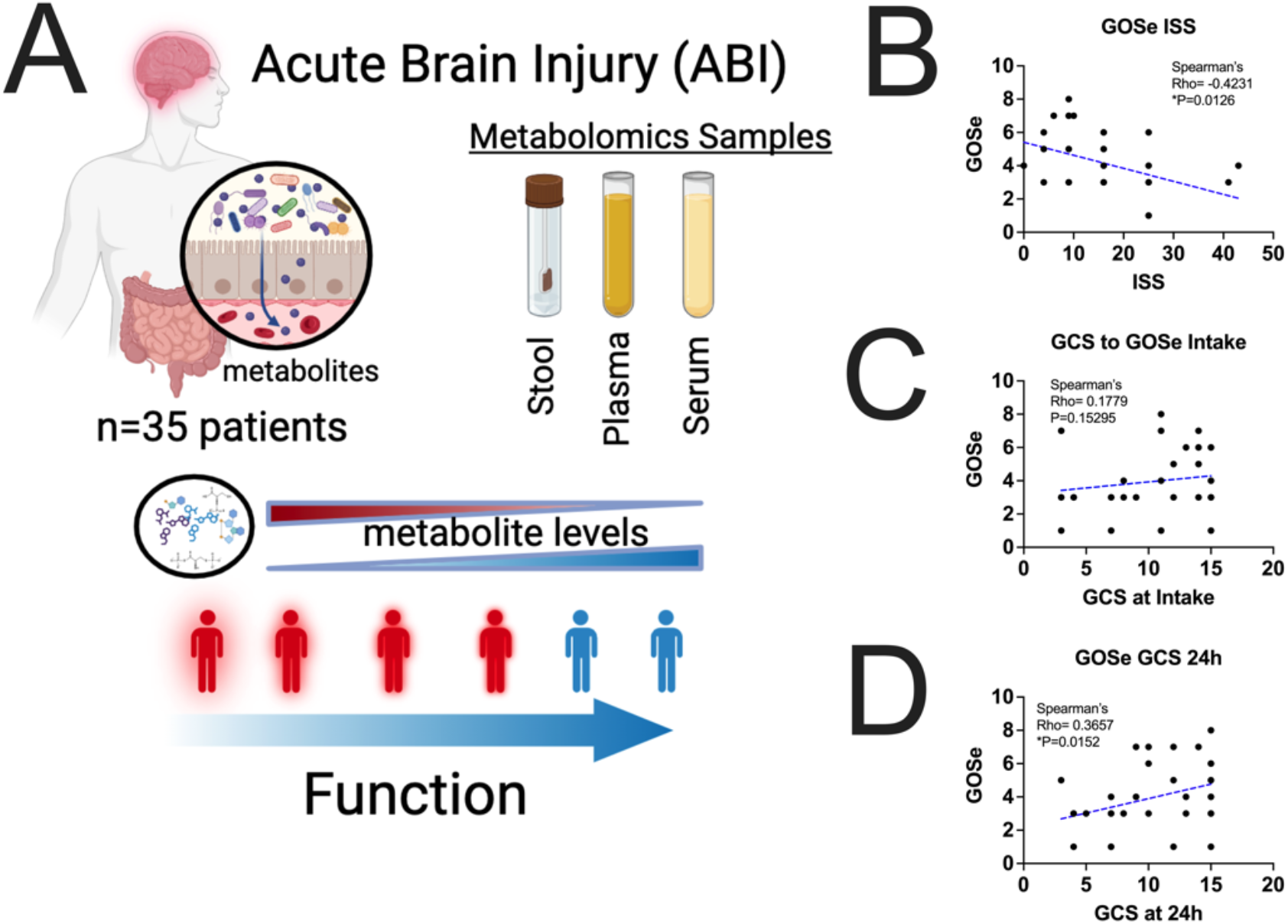
Study design. (**A**) Graphical study design. 35 patients with ABI were sampled cross-sectionally and longitudinally. 1 to 4 stool, serum, and plasma samples per patient were collected, with the earliest samples collected within 48h of injury and hospitalization. Targeted metabolomics of 300+ microbiome produced or modified metabolites were examined in stool, plasma, and serum with tandem shotgun metagenomics of stool and compared to the injury severity score (ISS) and Glasgow coma scale (GCS), GCS at 24h, and functional neurological outcome by the extended Glasgow outcome score (GOSe). (**B**) Association of ISS to GOSe was significant (Spearman, Rho= -0.4321, P=0.0151).(**C**) Association of GCS to GOSe was not significant (Spearman, Rho=0.1779, P=0.15296). (**D**) GOSe and GCS at 24h were significantly associated (Spearman, Rho=0.3657, *P=0.0152).

Inclusion criteria: (1) age ≥ 18 years; (2) admitted to the NCCU for ABI (including blunt and penetrating traumatic brain injury—TBI; intracranial hemorrhage—ICH; ischemic stroke—IS) with a GCS ≤ 12 or deemed to be high risk for deterioration based on brain imaging; (3) obtain consent within 48 hours from admission (either from the patient or surrogate decision maker).

Exclusion criteria: Patients with pregnancy, inflammatory bowel disease, normal brain imaging, pre-existing neurologic deficits, past neurologic or neurosurgical pathology. Moribund patients upon admission, patients with anticipated withdrawal of life-sustaining treatments (WOLST) in the first 72 hours of ICU admission, or patients with major comorbidities/organ failures besides the ABI were also excluded.

### Sample Collection

All human subjects in this study were consented and approved by the University of Chicago Institutional Review Board (IRB 22-0822). Fecal samples were collected by rectal swab or after bowel movements. Blood was collected with both heparin (+) and (-) tubes and centrifuged within 30 minutes (2000x*g*, 10min). All samples were frozen with N_2_(*l*) and stored at -80°C until processing ‘omic analysis.

### Metabolomics

Metabolomics were performed at the University of Chicago Duchossois Family Institute. Stool metabolomes were analyzed across four mass spectrometry platforms to capture quantitative and qualitative levels of metabolites with validation by retention time and fragmentation comparison to standards. Gas chromatography-mass spectrometry (GC-MS) was used for 222 compounds following derivatization with pentafluorobenzyl bromide (PFBBr) and trimethylsilyl-methoxamine (TMS-MOX) in two separate reactions. SCFAs (acetate, butyrate, propionate), lactate, and succinate were quantitated following PFB derivatization and negative collision induced-gas chromatography-mass spectrometry ((-)CI-GC-MS, Agilent 8890). Forty-eight PFB-derivatized compounds within SCFA, branched chain fatty acid, AA, aromatic, hydroxylated fatty acid, organic acid, and indole subclasses were studied by normalized peak area. Positive ion electron impact-GC-MS ((+)EI-GC-MS, Agilent 7890B) was used for 169 molecules in the organic acid, carbohydrate, TCA intermediate, sterol, AA, indole and fatty acid subclasses following TMS-MOX-derivatization. Negative mode liquid chromatography-electrospray ionization-quadrupole time-of-flight-MS ((-)LC-ESI-QTOF-MS, Agilent 6546) was used to analyze 49 BAs from the primary, secondary and glyco/tauro-conjugated subclasses were analyzed. Retention time validation, standard intact, and fragment masses were detected with differences < 5 ppm from calculated values. Positive mode LC-triple quadrupole-MS ((+)LC-ESI-QQQ-MS, Agilent 6547) was used for 34 indole and tryptophan catabolites. Downstream normalized values were corrected for batch effects using ‘*combat’* in R.

### Shotgun metagenomic sequencing

Sequences were filtered using Trimmomatic (v0.39) for PE (paired-end) reads, using default. Contigs for each sample were generated using megahit (v.1.2.9) and short reads mapped with samtools (v1.19.2), analyzed in anvi’o (v7.1). Open reading frames (ORFs) were identified using ‘*anvi-gen-contigs-database*’ and annotated for KEGG (Kyoto Encyclopedia for Genes and Genomes) Orthologs using ‘*anvi-run-kegg-kofams’*. Coverage tables were collapsed onto each KO as a sum of genes, and normalized by sample read count.

## RESULTS

Functional outcomes varied between patients (unfavorable outcome, GOSe 1 to 4, N=23; favorable outcome GOSe 5-8, N=12); 37.1% TBI, 20.0% ischemic stroke IS, 34.3% ICH, and 8.6% IS+ICH. Stool, plasma and serum were collected cross-sectionally and longitudinally (1 to 4 samples of each type collected per patient) with the earliest samples acquired within 48 hours of hospitalization (Supplementary Tables 1 and 2). Initial GCS was not correlated with outcome (Fig. 1, C, Spearman, P > 0.05). ISS was negatively correlated (Rs = -0.423, **P = 0.0126) and GCS at 24h was positively associated (Rs = 0.365, **P = 0.0152) (Fig. 1, B and D, Spearman). We then explored microbiome-metabolites associated with functional outcome.

### Microbiome-modified metabolites in serum and plasma are indicative of ABI outcome

Targeted metabolomics of 300+ metabolites produced or modified by the microbiome were measured in serum and plasma. These molecules spanned short chain fatty acids (SCFAs), bile acids (BAs), sugars, amino acids (AAs), indoles and tryptophan metabolites, and small aromatics. Metabolites not found in 50% of the subjects were removed to yield 100+ signals. To visualize the relationship between outcome after ABI and the serum and plasma metabolomes, we performed Principal Component Analysis using the first available samples (Fig. 2). Patients could be segregated between poor (GOSe 1-4, inset, red) and good outcomes (GOSe 5-8, inset, blue) with either serum or plasma, suggesting unique metabolite profiles after ABI (PERMANOVA, ***P<0.001). Segregation between outcomes could also be achieved using only the BAs (excluding all other metabolite classes) (Fig. 2, B, PERMANOVA, ***P<0.002), suggesting these microbiome-modified molecules are strongly associated with outcome.

**Figure 2.**
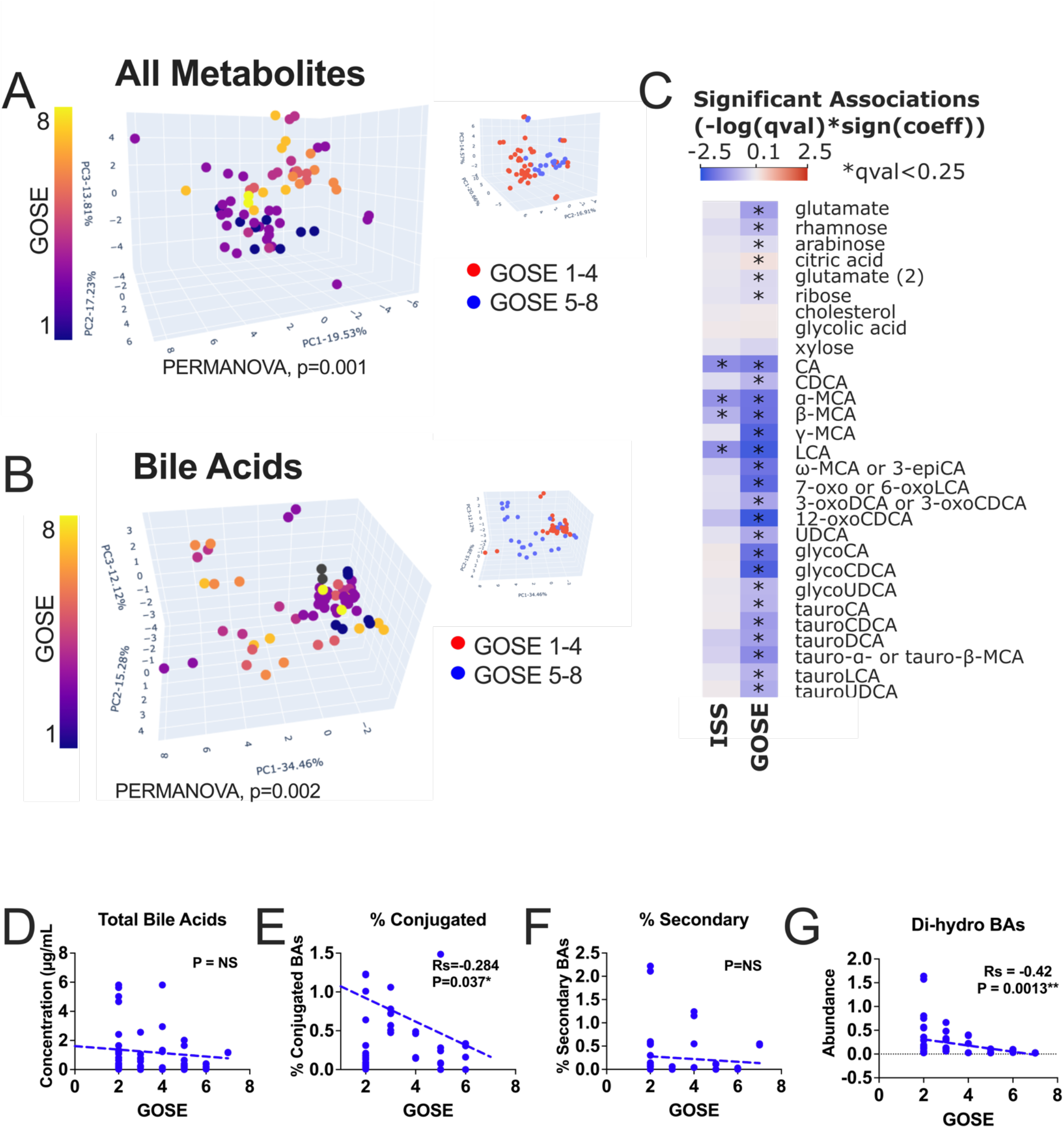
Microbiome-modified metabolites in serum and plasma are indicative of outcome after ABI. (**A**) Principal component analysis (PCA) reveals serum and plasma metabolites are distinct across outcomes (GOSe). Each dot is representative of the first serum or plasma sample and colored by GOSe score. Inset panel is colored by bad outcomes (GOSe 1-4, red) and good outcomes (GOSe 5-8, blue). Statistical significance was calculated for each GOSe group for each tissue type (serum or plasma, PERMANOVA, P=0.001). (**B**) Principal component analysis of bile acid signals found in serum and plasma. BAs are sufficient to discriminate across GOSe scores using the first plasma or serum sample, colored by GOSe score. Inst is colored by GOSe score 1-4 (red) or 5-8 (blue) and are statistically significant for either serum or plasma (PERMANOVA, P=0.002). (**C**) Significant associations (qval<0.25) for ISS and GOSe across all serum and plasma metabolites (-log(qval)*sign(coeff)) (MaAsLin2, red=positively associated, blue=negatively associated). More metabolites were predictive of GOSe than ISS. Model accounted for repeated measures, time of acquisition from intake, and tissue type. (**D-G**) Bile acid signals form the first sample acquired (within 48h of intake) such as percent conjugated (Spearman’s Rho = -0.284, *P=0.37) and di-hydro BAs (Spearman’s Rho = -0.42, **P=0.0013) are negatively associated while with outcome, while total BAs and secondary BAs are not (Spearman, two-tailed P>0.05)

To identify specific metabolites associated with outcome (GOSe), we performed multivariate analysis with MaAsLin2, a well-utilized algorithm for phenotype-multi-omic statistics using all serum and plasma samples, accounting for repeated measures, time of sample acquisition from hospital admission and tissue type (serum or plasma). After false discovery rate (FDR) correction, 26 metabolites were significantly associated with GOSe and 4 metabolites were associated with ISS, suggesting these signals are specific to brain injury (Fig. 2, C, *qval<0.25). Glutamate, a known mediator of neuronal excitotoxicity, was negatively correlated to GOSe, as were sugars rhamnose, arabinose and ribose. Citric acid was positively correlated with GOSe. Importantly, 20 BA species were negatively associated with GOSe, predominated by primary BAs cholic acid (CA) and chenodeoxycholic acid (CDCA), α,β, and γ-muricholate (MCA), and glycine and taurine conjugated BAs glyco-CA, glyco-CDCA, glyco-ursodeoxycholate (UDCA), tauro-CA, tauro-CDCA, tauro-deoxycholate acid (DCA), tauro-α/β-MCA, tauro-lithocholate (LCA) and tauro-UDCA.

The microbiome affects BA concentrations through deconjugation and secondary BA production. Using the earliest samples (within 48h), we found no significant association between GOSe and total BA levels or the percentage of secondary BAs (Fig. 2, D and F, Spearman, P>0.05), but percentage of conjugated BAs and di-hydroxyl BAs (CDCA, DCA and UDCA) were negatively associated (Fig. 2, E and G, Spearman, *P<0.05). Together, these data suggest specific microbiome-modulated metabolites including BAs are indicative of neurological homeostasis and highlights their potential in ABI diagnostics.

### The gut microbiome is associated with outcome

Levels of conjugated and primary BAs associated with outcome suggests impairment of microbiome function in the gut. Therefore, we directly examined the gut microbiome metabolome. Multivariate analysis accounting for repeated measures and time of sample collection yielded 24 unique metabolites significantly associated with outcome (Fig. 3, A, MaAsLin2, *qval<0.25). Twenty-one metabolites negatively associated with outcome included BA allocholate, uracil, psicose, AAs alanine, threonine, aspartate, glycine, isoleucine, proline, leucine, serine, phenylalanine, methionine, cysteine, and lysine, and fatty acids 5-aminovalerate and succinate. 10 positively associated metabolites included hydrocinnamate, *p*-cresol, sucrose, indole, palmitate, and myristic acid, cholesterol and its microbiome-converted product coprostanol, and BAs hydeoxycholate and α-muricholate.

**Figure 3.**
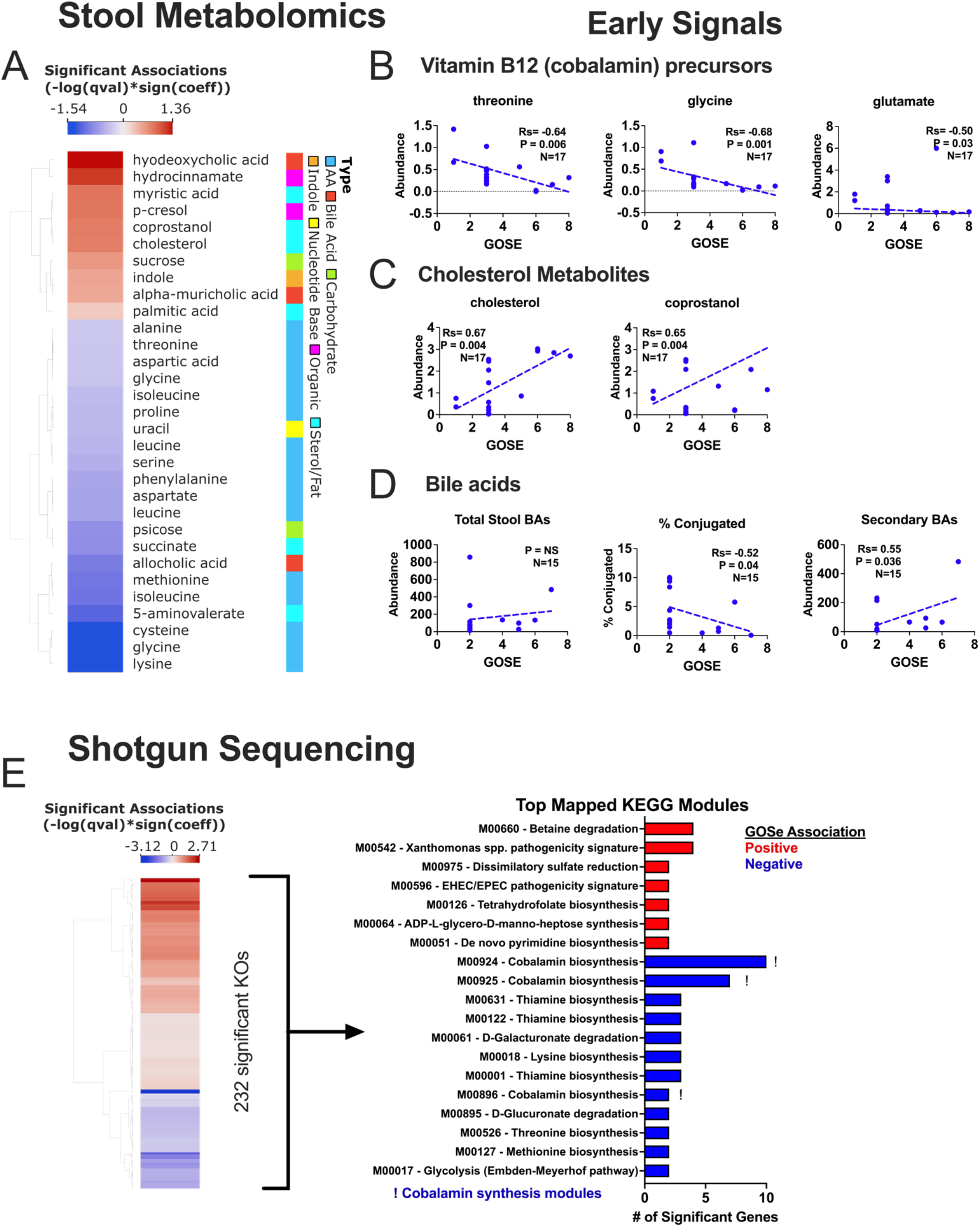
The microbiome is associated with outcome. **(A)** Significant associations between stool metabolites and GOSe (MaAslin2, *qval<0.25, red=positive associations, blue=negative associations). Model accounted for time of sample acquisition from intake and repeated measures. (**B**) Examination of the first samples provided (N=17) revealed a negative association between GOSe and the abundance of Vitamin B12 precursors threonine (Rs= -0.64), glycine (Rs=-0.68), and glutamine (Rs=-0.50) (Spearman, *P<0.04 for all correlations). (**C)** Positive associations were found between outcome and cholesterol (Rs=0.67) or coprostanol (Rs=0.65) (Spearman, *P<0.004). (**D)** Bile acid signals in stool were associated with outcome including a negative association with the percentage of conjugated BAs (Spearman’s Rho=-0.52, *P=0.04), a positive association with the abundance of secondary BAs (Spearman’s Rho=0.55, *P=0.036). Total bile acids in stool were no significantly associated (P>0.05, NS). (**E**) Microbes are enriched for vitamin B12 synthesis pathways. Shotgun sequencing of stool samples revealed 232 KEGG Orthologs (KOs) significantly associated with GOSe outcomes (MaAslin2, qval<0.02). Mapping of these KOs to KEGG molecules revealed that top modules (>1 significant KO mapping to that module) included vitamin B12 synthesis (M00924, M00925, and M00896).

The microbiome affects systemic levels of cholesterol and BAs by limiting available cholesterol and BA pools in the gut lumen in addition to effects of BA species on *de novo* BA synthesis via farsenoid X receptor (FXR). Furthermore, BAs and vitamin B12 (VB12) negatively influence each other’s absorption, with their levels being modified by gut microbiota. Given the circulating, outcome-associated BAs, we examined VB12 precursors, cholesterol, and BAs in the earliest available stool samples (within 48h). VB12 precursors threonine, glycine and glutamate were negatively associated with outcomes (Fig. 3, B, Spearman, *P<0.03 for all regressions). Conversely, cholesterol/coprostanol were positively correlated with outcomes (Fig. 3, C, Spearman, **P<0.004 for all regressions). Total BAs did not vary significantly with outcome, but percentage of conjugated (Spearman’s Rs=-0.52) and secondary BAs (Spearman’s Rs=0.55) were negatively and positively associated, respectively (Fig. 3, D, *P<0.04).

To further explore the microbiome, we performed shotgun sequencing to measure the microbiota functional potential. We identified 232 KOs (Kyoto Encyclopedia of Genes and Genomes Orthologs) associated with outcome (Fig. 3, E). These KOs mapped to 19 modules; grouped enzymatic reactions. Of these, cobalamin biosynthesis modules (M00924, M00925 and M00896) were negatively associated revealing that microbes capable of VB12 synthesis bloomed in patients with adverse outcomes. Together, these data implicate VB12 and cholesterol balance with concomitant microbiome processes as markers of brain injury outcome.

### Network of metabolomic features across tissues identifies tauro-α/β-muricholic acid as central to outcome

To examine metabolite relationships across tissues, we generated a metabolite network with outcome (primary metabolite-to-outcome associations), and each other (secondary metabolite-to-metabolite associations) (Fig. 4, A). The network was built using the earliest time-matched stool, plasma, and serum samples (N=17 patients) with P<0.05 and P<0.01 cutoffs for primary and secondary associations, respectively. The final network consisted of 127 metabolites connected by 304 edges, with 4.79 average neighbors. Generally, features from the same tissue covary, e.g. stool features covary with other stool metabolites. However, tissue types covary for negatively associated metabolites (Fig. 4, A, Cluster II) more than positively associated metabolites (Cluster I) suggesting greater crosstalk between the gut and systemic circulation for metabolites that negatively affect outcome.

**Figure 4.**
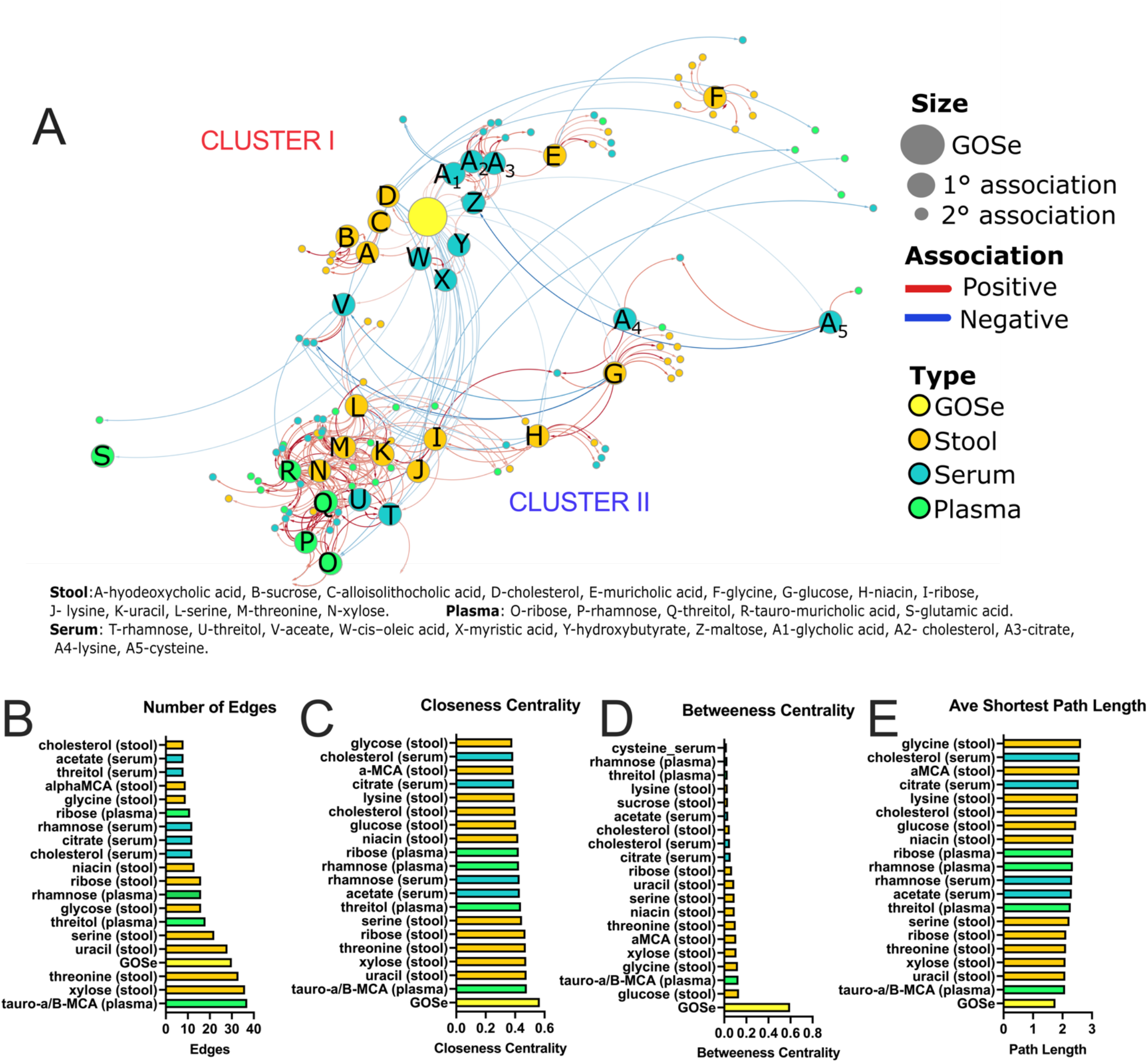
Cross-tissue metabolite networks. **(A)** Network analysis of tissue, plasma, and stool metabolites associated with outcome (GOSe). Metabolites with a significant 1° association with GOSe (Spearman, P<0.05), or 2° association (P<0.01) are represented as nodes colored by tissue of origin (stool=orange, serum=blue, plasma=green). 1° positively associated metabolites are grouped in Cluster I with 1° negatively associated metabolites in Cluster II. Edges are colored by positive (red) or negative (blue) associations. **Stool**:A-hyodeoxycholic acid, B-sucrose, C-alloisolithocholic acid, D-cholesterol, E-muricholic acid, F-glycine, G-glucose, H-niacin, I-ribose, J-lysine, K-uracil, L-serine, M-threonine, N-xylose. **Plasma**: O-ribose, P-rhamnose, Q-threitol, R-tauro-muricholic acid, S-glutamic acid. **Serum**: T-rhamnose, U-threitol, V-acetate, W-*cis–*oleic acid, X-myristic acid, Y-hydroxybutyrate, Z-maltose, A1-glycholic acid, A2-cholesterol, A3-citrate, A4-lysine, A5-cysteine. **(B-E)** Top 20 node of importance characterized by connectivity (number of edges, **B**), closeness centrality (**C**), betweenness centrality (**D**), and average shortest path length (**E**).

Network-based analyses enable inference of keystone features that underpin the network by examining placement and connectivity. We used 4 metrics: number of edges, closeness centrality, betweenness centrality, and average shortest path length (Fig. 4, B-E, respectively). Of these features, we found the top 20 features were consistent across all 4 metrics including plasma tauro-α/μ-MCA, stool xylose and glucose, and stool threonine, highlighting the potential importance of these features in brain injury outcome. Interestingly, tauro-α/μ-MCA is a naturally occurring FXR-antagonist, and could contribute to dysregulation of cholesterol homeostasis during ABI.

## Discussion

We examined microbial metabolites to identify molecules predictive of outcome in patients with acute stroke and traumatic brain injury admitted to a neurosciences critical care unit. We identified a class of microbiome metabolites, bile acids, associated with functional outcomes at hospital discharge. While no significant changes in total BA concentrations were detected, shifts towards conjugated and hydrophilic BAs were indicative of worse outcomes including tauro-α/μ-muricholate, tauro-ursodeoxycholate, and glyco-ursodeoxycholate. This is particularly relevant considering the role of BAs in regulating metabolic-homeostasis as ligands of the energy integrator FXR.^14^ While most BAs are FXR-agonists, TMCA, TUDCA, and GUDCA are FXR-antagonists.^15^ BAs are deconjugated by the gut microbiome. Studies in TBI patients have shown increased conjugated species (taurocholate, glycocholate) and suggested a link between BA accumulation (secondary to microbial dysbiosis or splanchnic ischemia) and TBI induced platelet dysfunction.^16^ Further putative mechanisms involve blood-brain barrier (BBB) disruption, allowing BAs to readily enter the brain wherein they may alter homeostasis.^17,18^ BA metabolism has been correlated with outcome in neurodegenerative disorders with significantly lower serum concentrations of a primary BA (cholate [CA]) and increased levels of bacterially produced, secondary BAs, deoxycholate and its glycine/taurine forms. Higher levels of secondary conjugated BAs (GDCA, GLCA, and TLCA) were associated with worse cognitive function. In follow-up studies, these changes correlated with alterations in cerebrospinal fluid disease markers and brain imaging.

Cell lines, animal, and human studies suggest secondary BAs, particularly DCA, disrupt mitochondrial membranes to affect reactive oxygen species, inflammation, and apoptosis while decreasing cell viability and DNA synthesis. Studies in rodents demonstrated deuterium-labeled DCA crosses the BBB and increases its permeability^19,20^. Secondary BAs in blood may enter the brain through BBB permeability, affecting brain physiology and metabolism^21^ with the entire complement of BAs detected in the brain.^15,19,22,23^ It remains unclear whether transport from the periphery, local synthesis, or both is responsible. The function of BAs in the brain is undeciphered, with support for neurosteroid effects^24–26^. Preclinical and clinical evidence indicates that acute cerebral ischemia causes intestinal dysfunction, affecting gut dysbiosis, which then drives pathogenesis^27–30^.

Brain injury patients may exhibit increased blood glucose and cholesterol, with the latter implicated in causing brain inflammation and secondary damage^19,23^. Hepatic FXR-deletion increases circulating cholesterol and glucose levels, impairing cholesterol efflux while FXR-agonism increases efflux.^20^ We detected decreased cholesterol in stool consistent with FXR-antagonism, reducing efflux and absorption. This finding highlights the need for further research on BA regulation of FXR during ABI. Understanding effects of specific BAs on the brain may yield insight into SBI and potentially unify previously reported molecular and clinical phenotypes.

There is a critical need to identify biomarkers that reflect early transitions to dysmetabolism, particularly those that are causal and modifiable. This study underscores the importance of the microbiome in modifying key molecules, with detection within 48h suggesting that these events occur early. However, when dysbiosis alters BAs, systemic and neurological effects can occur, though the molecular triggers for metabolomic shifts remain unknown. Lastly, individual-specific microbiomes may have a greater predisposition to produce molecules that potentiate SBI.

Pre-clinical animal models of brain injury are required to demonstrate the significance of these microbiome metabolites as drivers of SBI. These models may deconvolve the series of events after brain injury and inform novel interventional strategies. Brain injury models of mice can be combined with standard microbiome models; Germ-free mice, fecal material transplants, and direct administration of BAs and FXR chemical modulators.

## Data Availability

All shotgun sequencing data was submitted to the sequence read archive (SRA). All derived metabolomics data was deposited to the National Metabolomics Data Repository (NMDR) and can be found at Metabolomicsworkbench.org. All raw metabolomics data was submitted to MassIVE. All deidentified sample identifiers and relevant metadata was submitted and can be freely downloaded with both sequencing and metabolomics data.

## Abbreviations

TBI: traumatic brain injury
ABI: acute brain injury
SBI: secondary brain injury
GCS: Glasgow Coma Scale
ISS: Injury Severity Score
GOSe: Glasgow Outcome Scale extended

## Acknowledgments

We thank the Duchossois Family Institute (DFI) at the University of Chicago for their generous funding. We thank the nurses, training physicians, and attendings of the University of Chicago neurocritical care unit for the invaluable help in completing this project.

## Author Contributions

CL, EC, and OD conceptualized and designed the study. OD and HC analyzed the data. CL, EC, OD, HC, WR and AS interpreted the data. SR, NB, WR, DB and CL consented patients and collected the samples. OD, HC and AS prepared the samples. AS performed the mass spectroscopy. EC and CL supervised the study. OD and CL wrote the manuscript. All authors contributed to the editing of the manuscript.

