## Supplementary figures and images for "MICROBIOME-MODIFIED METABOLITES PREDICT ACUTE BRAIN INJURY OUTCOME"

### Supplemental Table 1

**Supplementary Table 1 – Patient Characteristics**

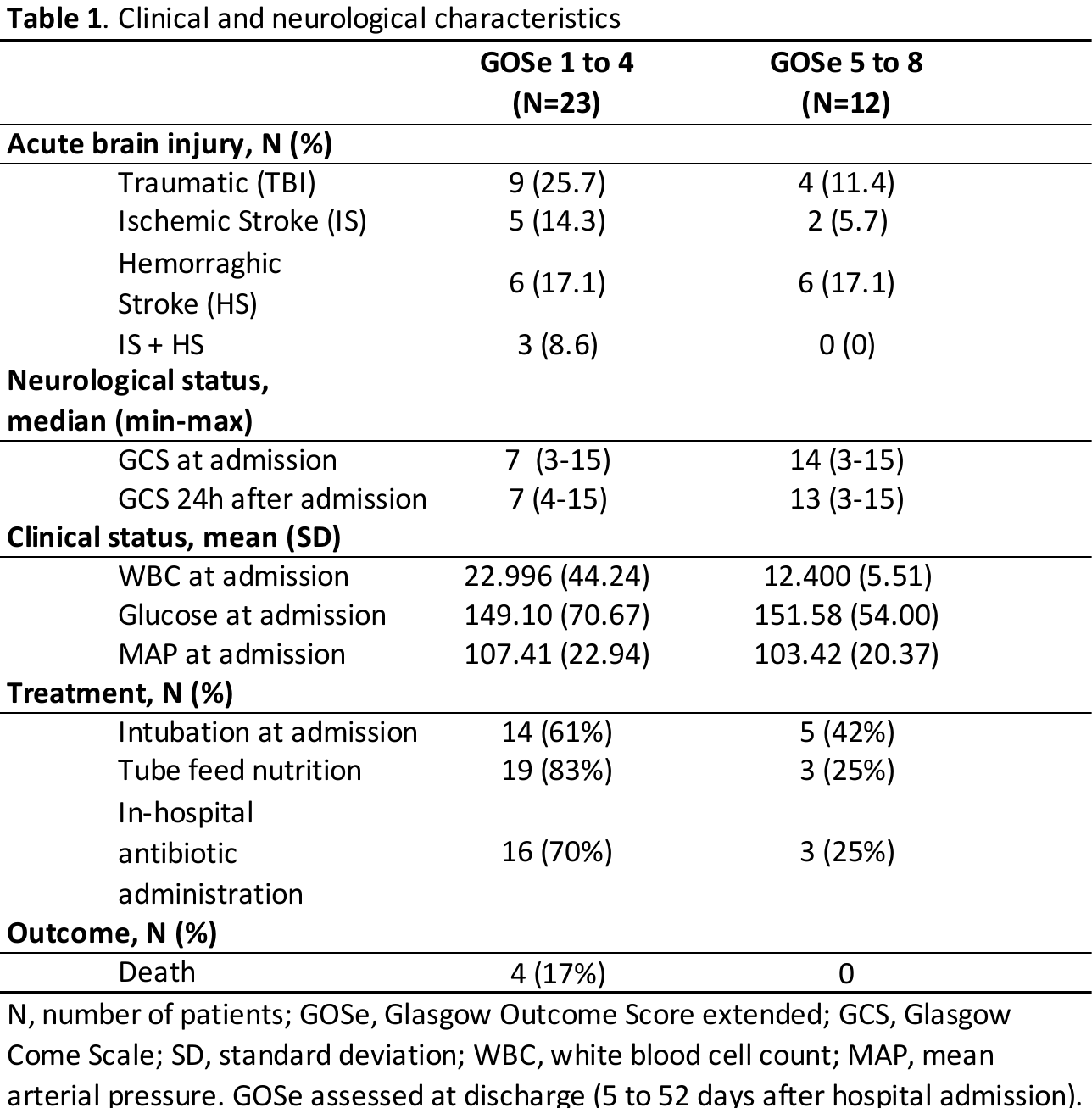
