## Supplemental Table 2 for "MICROBIOME-MODIFIED METABOLITES PREDICT ACUTE BRAIN INJURY OUTCOME"

|  | Diagnosis | Description of CT on admission | GCS at presentation | Injury Severity Scale (ISS) | Duration of ICU stay | Duration of hospital stay |
| --- | --- | --- | --- | --- | --- | --- |
| 1 | Penetrating TBI (GSW) | R Frontoparietal IPH and SAH  R parietal SDH  No MLS  Patent basal cistern | 11 | 9 | 5 | 16 |
| 2 | Ischemic stroke | R frontoparietal cytotoxic edema and loss od grey white matter differentiation  No midline shift  Basal cistern patent | 14 | 4 | 7 | 19 |
| 3 | Blunt TBI (MVA) | Bilateral diffuse axonal injury  No MLS  Patent basal cistern  L ICA dissection | 4 | 16 | 24 | 24 |
| 4 | Hemorrhagic stroke (Hypertensive) | L thalamic IPH  Extension to L midbrain and pons  Intraventricular extension  No MLS  Patent basal cistern | 3 | 25 | 8 | 8 |
| 5 | Ischemic stroke with hemorrhagic conversion | R frontal IPH  Vasogenic edema edema  Intraventricular extension  11 mm MLS  Patent basal cistern | 15 | 9 | 3 | 42 |
| 6 | Hemorrhagic stroke (Hypertensive) | R basal ganglia IPH with surrounding edema  Intraventricular extension  5 mm MLS  Patent basal cistern | 15 | 9 | 3 | 28 |
| 7 | Blunt TBI (MVA) | R SDH  L Frontoparietal IPH and contusion  12 mm MLS  Effaced basal cistern  Frontal bone and craniofacial fractures  Unruptured L cavernous ICA aneurysm | 3 | 41 | 10 | 34 |
| 8 | Hemorrhagic stroke (Hypertensive) and ischemic stroke | R basal ganglia IPH with surrounding edema  5 mm MLS  L ventral pontine infarct  Patent basal cistern | 7 | 25 | 25 | 25 |
| 9 | Hemorrhagic stroke (Hypertensive) | R basal ganglia IPH  No MLS  Patent basal cistern | 7 | 9 | 7 | 14 |
| 10 | Ischemic Stroke | No evidence of ischemic or hemorrhagic changes on CT  (Subsequent MRI with frontotemporal diffusion restriction)  No midline shift  Basal cistern patent | 15 | 4 | 2 | 7 |
| 11 | Ischemic stroke | Perfusion imaging with core infarct of L MCA territories  Associated hemorrhagic conversion and intraventricular extension  4.5 mm MLS  Effaced basal cistern | 11 | 25 | 12 | 12 |
| 12 | Hemorrhagic stroke (Hypertensive) | Thalamic IPH with intraventricular extension  No MLS  Patent basal cistern | 7 | 9 | 17 | 33 |
| 13 | Hemorrhagic stroke (cerebral amyloid angiopathy) | R parietal and L occipital IPH  No MLS  Patent basal cistern | 14 | 9 | 3 | 10 |
| 14 | Hemorrhagic stroke | L basal ganglaia, L temporo  Frontoparietal IPH with associated edema  L frontoparietal SAH  12 mm MLS  Uncal herniation | 11 | 25 | 3 | 8 |
| 15 | PBI (GSW) | R temporal, frontoparietal and occipital hematomas with associated edema  R SDH  13 mm MLS  Effaced basal cistern  Nonocclusive traumatic thrombosis of sagittal sinus and R transverse sinus | 4 | 25 | 38 | 52 |
| 16 | Ischemic stroke | L frontoparietal, temporal and occipital loss of grey white matter differentiation and hypoattenuation  10 mm MLS  Effaced basal cistern | 11 | 9 | 5 | 21 |
| 17 | Blunt TBI (MVA) | L frontaopareital and temporal IPH and Bl SDH  3 mm MLS  Basal cistern patent  Frontal, sphenoid, L parietal and L temporal bones fractures | 4 | 16 | 18 | 31 |
| 18 | Blunt TBI (MVA) | Bl frontotemporal contusion and IPH. R occipital and cerebellar epidural hematomas, bilateral frontotemporal contusions. bl SDH  5 mm MLS  R transverse sinus thrombosis  R temporal bone fracture | 15 | 16 | 23 | 32 |
| 19 | Hemorrhagic stroke (Hypertensive) | L basal ganglia IPH  4 mm MLS  Patent basal cistern | 15 | 0 | 6 | 11 |
| 20 | Blunt TBI (fall) | Bifrontal and R cerebellar IPH and contusion. BL SAH and SDH  4 mm MLS  Patent basal cistern  Occipital and orbital fractures | 8 | 25 | 5 | 5 |
| 21 | Penetrating TBI (GSW) | L Frontoparietal and, temporal IPH and SAH with surrounding edema  Bl SDH  7 mm MLS  Haeorrgae into basal cistern  No significant brainstem injury  Left maxillary and ethmoid hemosinus from displaced fractures | 3 | 16 | 19 | 23 |
| 22 | Ischemic Stroke | L frontoparietal and temporal hypoattenuation and loss of grey white matter differentiation  Surrounding cytotoxic edema  6 mm MLS  Effaced basal cistern  R MCA occlusion | 14 | 25 | 28 | 28 |
| 23 | Ischemic Stroke | L frontoparietal, temporal and occipital hypoattenuation and loss of grey white matter differentiation  Surrounding cytotoxic edema  8 mm MLS  basal cistern patent  No significant brainstem injury  R MCA occlusion | 15 | 25 | 4 | 12 |
| 24 | Hemorrhagic stroke (Hypertensive) | R thalamic IPH with intraventricular extension and surrounding edema  No midline shift  Basal cistern patent | 14 | 9 | 12 | 34 |
| 25 | Hemorrhagic stroke (Hypertensive) | L basal ganglia IPH with surrounding edema  3 mm midline shift  Basal cistern patent | 12 | 16 | 1 | 4 |
| 26 | Ischemic Stroke | Frontotemporal loss of grey white matter differentiation  6 mm midline shift  Basal cistern patent  R MCA occlusion | 9 | 16 | 21 | 21 |
| 27 | Blunt TBI (fall) | R SDH  L lateral ventricle IVH  8 mm MLS  Basal cistern patent | 11 | 16 | 14 | 22 |
| 28 | Blunt TBI (MVA) | Bl frontotemporal IPH and contusions with surrounding edema  No MLS  Basal cistern patent | 3 | 10 | 6 | 10 |
| 29 | Hemorrhagic stroke (Hypertensive) | L thalamic IPH with intraventricular extension  No MLS  Basal cistern patent | 15 | 4 | 2 | 5 |
| 30 | Blunt TBI (fall) | L SDH  L SAH  5 mm MLS  Basal cistern patent  L temporal Contusion  R temporal bone fractur | 13 | 25 | 11 | 21 |
| 31 | Blunt TBI (fall) | Bl Scalp hematomas  No acute intracranial hemorrhage  No MLS  Basal cistern patent  Incidental 7 mm basilar tip aneurysm, unruptured | 3 | 6 | 2 | 5 |
| 32 | Hemorrhagic stroke (Hypertensive) | R basal ganglia IPH  No MLS  Basal cistern patent | 15 | 16 | 2 | 11 |
| 33 | Blunt TBI (MVA) | L epidural hematoma overlying temporal lobe  L temporal fracture  R temporal contusion and SAH  R and falcine SDh  SAH in basal cistern  Uncal herniation | 8 | 43 | 7 | 16 |
| 34 | Hemorrhagic stroke (Hypertensive) | R thalamic IPH  5 mm MLS  Basal cistern patent | 14 | 16 | 4 | 12 |
| 35 | Ischemic stroke with hemorrhagic conversion | R basal ganglia IPH with intraventricular extension  4 mm MLS  Basal cistern patent | 12 | 25 | 6 | 25 |
